# FLiPA: A versatile platform for quantitative analysis of protein-glycosphingolipid interactions

**DOI:** 10.64898/2026.07.13.738194

**Authors:** Shannon J. McKie, Ella Bishop, Janet E. Deane

## Abstract

Interactions between proteins and glycosphingolipids (GSLs) regulate various cellular processes and altered GSL metabolism contributes to numerous diseases. The diverse glycan headgroups and ceramide backbones of GSLs shape membrane organisation, fluidity, curvature, and tension. As protein recognition frequently depends on both glycan specificity and the organisation of GSLs within the membrane, these interactions remain challenging to characterise in vitro. Here, we introduce FLiPA (Fluorescent Liposome Plate Assay), a versatile method that utilises fluorescent agarose-embedded giant liposomes for the quantitative analysis of protein-GSL interactions. By enabling systematic control of membrane and buffer composition, FLiPA provides an accessible and robust platform for dissecting the molecular determinants of protein-GSL interactions, including the roles of cholesterol, membrane order, protein oligomerisation and ionic strength.

## Introduction

Protein-lipid interactions are an essential component of many cellular processes and disruption of these interactions is implicated in the pathogenic mechanism of numerous diseases. Therefore, understanding them advances fundamental molecular understanding and also guides research on the development of novel treatment strategies. However, robust and accessible methodologies for the *in vitro* characterisation of protein-lipid interactions remain limited. This is especially relevant for a class of lipid called the glycosphingolipids (GSLs), which are formed from a ceramide backbone and a glycan headgroup (1). There is significant variability in the molecular structure of both the ceramide and glycan portions, giving rise to a compositionally diverse lipid class from which specific protein-GSL interactions can arise (2). GSLs are enriched in the outer leaflet of the plasma membrane (PM) where they play key roles in cell signalling, ion channel modulation and cell-cell adhesion (3). These lipids are particularly abundant in the nervous and immune systems and are exploited by pathogens for host-cell attachment and invasion (4-6). Disruption of GSL metabolism is implicated in the pathogenesis of multiple neurodegenerative disorders, autoimmune diseases and cancers (7-12). Knowing the key determinants of specificity for protein-GSL interactions is therefore crucial to understanding how these vital lipids support cell development and function.

Characterisation of protein-GSL interactions support that the glycan headgroup helps to confer specificity (13). However, many monovalent protein-carbohydrate interactions are notoriously weak which can hinder biochemical quantification (14). In biology, glycan-mediated interactions often rely on multivalency to enhance binding avidity and enable physiologically relevant functions (15). Multivalency is achieved by clustering carbohydrate substrates and/or combining multiple carbohydrate binding domains of proteins through oligomerisation or membrane clustering. This facilitates multiple simultaneous interactions, which transforms the low-affinity monovalent interaction into a functional high-avidity interaction (16). Some of the most thoroughly characterised multivalent protein-GSL interactions come from viruses. For example, the capsid of Simian virus 40 (SV40) is formed from 72 pentamers of the VP1 protein, with each VP1 binding a single glycosphingolipid, specifically GM1. The monomeric Kd for the VP1-GM1 interaction is 1-5 mM, however the avidity of the interaction is enhanced by the formation of VP1 pentamer, and furthermore by constructing the viral capsid, enabling SV40 host cell entry (17, 18). Similar mechanisms are used by a number of other viral proteins including cholera toxin B protein binding to GM1 and Shiga toxin B-subunit binding to Gb3, and in all 3 viral examples, loss of the GSL receptor on host cells results in resistance to infection (18-20). In humans, myelin-associated glycoprotein (MAG, Siglec-4), crucial for signalling between oligodendrocytes and neurones, engages GSLs through a multivalent mechanism involving neuronal GSL clustering and oligodendrocyte-bound MAG homodimerisation (21, 22).

Characterising GSL-protein interactions *in vitro* is further complicated by the significant influence that GSLs exert on membrane architecture, including lateral organisation, curvature, fluidity and tension. GSLs are structurally and biochemically distinct from phospholipids with a strong tendency to laterally segregate into ordered GSL-rich domains known as lipid rafts or membrane microdomains, forming platforms that support GSL-dependent mechanisms (23). The molecular basis for the lateral segregation of GSLs is driven by cooperative interactions between GSL molecules, particularly the formation of a hydrogen bond network that arises due to the amide, carboxyl and carbonyl groups within the ceramide. In addition, the hydrophilic and bulky glycan headgroups exhibit variable hydration, charge and curvature properties, dependent on the specific glycan structure, that also influence lateral organisation (24). To balance these molecular signatures, GSLs require precisely controlled lipid environments to prevent lipid aggregation and membrane collapse, allowing the headgroups to be appropriately displayed on the surface of the lipid bilayer (24, 25).

Liposome-based approaches have been pivotal membrane models in the characterisation of protein-lipid interactions, revealing key insights that have expanded our understanding of how cells utilise these two distinct molecules to transmit information (26). A major advance in methodologies for exploring protein-lipid interactions was the development of the Liposome Micro Array (LiMA), which demonstrated the use of fluorescent agarose-fixed giant liposomes for high-throughput screening, facilitating specificity determination for a number of GFP-fused lipid-binding proteins (27, 28). However, a substantial gap remains in robust methodologies for expanding our understanding of protein-GSL recognition, particularly regarding specificity and membrane organisation. Here, we present FLiPA (Fluorescent Liposome Plate Assay), a versatile platform that combines gel-assisted giant liposome formation with fluorescence microscopy to quantitatively interrogate and visualise protein-GSL interactions. FLiPA utilises a 96-well plate format, affording medium throughput capacity and does not require additional specialist equipment enabling implementation across the research community. Using FLiPA, we show that protein-GSL interactions are highly sensitive to lipid composition, bilayer organisation, ionic strength, and protein oligomerisation state. These results demonstrate a significant advancement in the development of tools to explore protein-GSL interactions, establishing FLiPA as a powerful framework for the mechanistic study of these crucial molecular contacts.

## Results

### FLiPA principles and optimisation of GSL incorporation

The principles of FLiPA (Fig. 1a) begin with forming a thin agarose layer (TAL) at the bottom of each well of a chloroform-resistant flat-bottom 96-well plate using 1% ultra-low gelling type IX agarose. A lipid mixture of defined composition containing 0.2-0.5% dioleoyl phosphatidylethanolamine fluorescently tagged with Liss Rhodamine (Rhod-DOPE), is dissolved in a chloroform-based solvent and pipetted onto the TAL and left until the solvent has fully evaporated. Giant liposomes are formed in the TAL layer by the addition of hydration buffer containing the fluorescently-tagged protein of interest (POI). The TAL enables tethering of the liposomes to facilitate effective visualisation by microscopy. The liposomes are weakly tethered to the surface and excessive pipetting results in liposome loss from the surface. Including the POI in the hydration buffer overcomes this problem and we have not observed non-specific binding to carrier lipids. An alternative use of this method (Fig. 1a, part 3ii) is to hydrate the liposomes in the plate, agitate the plate to dislodge the liposomes into suspension and visualise them using a glass or cavity slide. This allows the addition of protein and other additives after the liposomes are hydrated as some may impact liposome hydration, and the glass/cavity slide format decreases background GFP fluorescence but also throughput. The fluorescent bioreporters utilised here possess an N-terminal his-tag and a C-terminal GFP fused to the soluble GSL-binding domains of siglec-7, galectin-3 and neurofascin isoform 155 (NF155) (Fig. 1b, Supplementary Table 1). Recruitment of protein to the membrane is monitored using fluorescence microscopy either as TAL-tethered layers or suspended in solution (Fig. 1c). Microscopy images are analysed to quantify protein-GSL interactions using a similar approach to the work pioneered in LiMA (27, 28). Specifically, the intensity of protein fluorescence is measured in the regions that colocalise with lipid to calculate the normalised binding intensity (NBI, ratio of GFP to Rhod-DOPE fluorescence) (Supplementary Fig. 1a).

**Figure 1:**
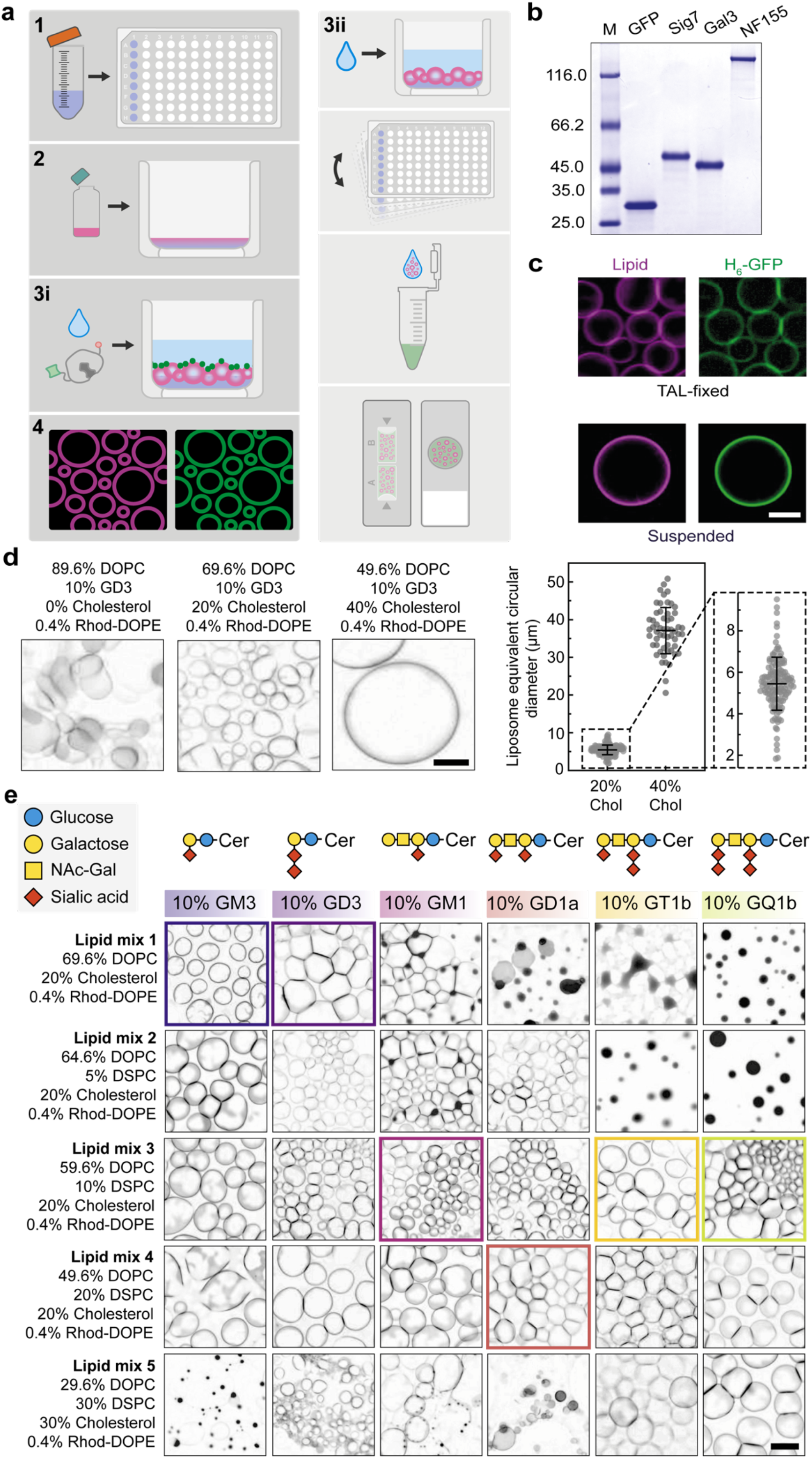
Fluorescent liposome plate assay (FLiPA) protocol overview and optimisation. **(a)** FLiPA workflow. (1) Each well of a chloroform-resistant 96-well plate is coated with a thin agarose layer (TAL). (2) 9 nmol of lipid mixtures in the appropriate organic solvent are added to the TAL. (3a) The liposomes are hydrated in buffer containing 0.25-0.5 µM GFP-tagged protein. (3i) Alternatively, hydrate the liposomes in buffer, agitate the plate to suspend the liposomes, transfer the liposomes to a tube containing GFP-tagged protein and any other additives, and then transfer to a cavity/glass slide. (4) Finally, image the plate or cavity/glass slide using fluorescence microscopy. **(b)** Coomassie-stained SDS-PAGE of the GFP-fused proteins utilised. GFP is H_6_-GFP, sig7 is the V-set domain of human siglec-7 with an N-terminal H_10_ tag and a C-terminal sfGFP tag, Gal3 is the CRD of human galectin-3 with an N-terminal H_6_ tag and a C-terminal eGFP tag, and NF155 is the full extracellular domain of neurofascin isoform 155 with an N-terminal H_10_ tag and a C-terminal sfGFP tag. **(c)** Liposome example images for both TAL-fixed in a plate and suspended in a cavity slide. Liposome composition was 79.5% DOPC, 10% NTA(Ni)-DGS, 10% cholesterol, 0.5% Rhod-DOPE, and they were hydrated in 25 mM Tris pH 7.4 and 150 mM NaCl containing 0.1 µM H_6_-GFP. Scale bar is 20 µm and images taken at 20x magnification. **(d)** Modulation of formation and size of liposomes containing 10% GD3 with increasing mol% cholesterol. Scale bar is 20 µm and images taken at 20x magnification. Liposome equivalent circular diameter is plotted for 20 (no. liposomes = 131) and 40 (no. liposomes = 56) mol% cholesterol ± S.D. **(e)** Optimisation of GSL incorporation for 6 GSLs, GM3, GD3, GM1, GD1a, GT1b and GQ1b by varying DOPC:DSPC ratio and cholesterol mol%, hydrated in 25 mM Tris HCl. Coloured boxes indicate liposome mixture selected for each GSL for GSL bioreporter screening. Scale bar is 50 µm and images taken at 10x magnification.

Related approaches have primarily focussed on the analysis of protein interactions with phospholipids and have not demonstrated their application to GSLs. The chemical and structural properties of GSLs make them a particularly challenging lipid class for giant liposome formation. GSLs have saturated ceramide backbones and large, often negatively charged, carbohydrate headgroups with high conformational flexibility (29). They are also able to form intermolecular hydrogen bonds and have high melting temperatures. Consequently, GSLs substantially influence membrane tension, fluidity and curvature and mix poorly with unsaturated carrier lipids leading to lipid aggregation and membrane collapse. This prevents the head groups from being arrayed correctly and compromises protein-GSL binding. Therefore, it was essential that bespoke lipid mixtures were optimised for each of the 6 GSLs explored here to promote consistent incorporation of each GSL. For the simpler, unbranched GSLs, GM3 and GD3, the only addition required to form giant liposomes, besides the standard unsaturated carrier lipid dioleoyl phosphatidylcholine (DOPC) and 0.2-0.5 mol% Rhod-DOPE, is cholesterol. Without cholesterol, the GSL mixtures fail to hydrate correctly, with some regions aggregating or forming dense, tubular membrane structures. The addition of 20% cholesterol yielded well-formed liposomes with a mean diameter of 5.5 µm ± 1.3 µm (± SD) while 40% cholesterol produced liposomes with a mean equivalent circular diameter of 37.1 µm ± 6.1 µm (± SD), demonstrating that varying cholesterol % can modulate liposome size (Fig. 1d, supplementary Fig. 1b). This feature is important for different applications of FLiPA, depending on which cellular membrane is being modelled or if the POI senses membrane curvature.

The addition of cholesterol alone was not sufficient for the reliable incorporation of the other 4 GSLs tested: GM1, GD1a, GT1b and GQ1b. These GSLs have larger, branched headgroups which required counterbalancing with the addition of saturated lipid to increase membrane ordering and packing. This leads to better toleration of the bulky and charged GSL headgroup and optimal ceramide packing. Here we used distearoyl phosphatidylcholine (DSPC) as an additional saturated carrier lipid so the headgroup presentation remained consistent even when the ratio of DOPC:DSPC changed. Sphingomyelin, also with a phosphocholine headgroup, was less effective at improving GSL incorporation than DSPC in this study. We varied the mol% of DSPC from 0-30% and cholesterol from 20-30% and identified mixtures for each GSL that consistently yielded well-formed giant liposomes (Fig. 1e). GSLs with larger headgroups required more DSPC to stabilise the membrane but a variable range of lipid mixtures was found to be effective for each GSL. This systematic analysis identified that lipid mix 3 (59.6% DOPC, 10% DSPC, 20% cholesterol, 0.4% Rhod-DOPE) supported liposome formation for all GSLs at 10% concentration. However, over time GM3 and GD3 were less stable in lipid mix 3 than in lipid mix 1 or 2. Therefore, for further analysis we selected the most reliable mix for each individual lipid: lipid mix 1 for GM3 and GD3; lipid mix 3 for GM1, GT1b and GQ1b; and lipid mix 4 for GD1a. While lipid mix 4 with 10% GT1b formed stable and numerous liposomes, it was not selected to explore protein binding due to lipid phase separation reducing GFP and Rhod-DOPE colocalization, which compromised the NBI quantification. Through optimisation of membrane composition, we have demonstrated the importance of the presence of cholesterol and an ordered lipid species for the reliable incorporation of 6 GSLs, providing a valuable framework for inclusion of additional GSLs in the future.

### FLiPA reveals GSL binding specificity and affinity of GFP-fused bioreporters

To demonstrate the effectiveness of FLiPA for quantifying protein binding to GSLs we selected three proteins with robust evidence of GSL interactions: siglec-7, galectin-3 and NF155. The GSL-binding domain of each was cloned into a vector with an N-terminal His-tag for purification and a C-terminal GFP for visualisation. The first bioreporter, siglec-7 is a member of the sialic acid binding immunoglobulin-like lectin (Siglec) family, and functions as an immune response inhibitor on the surface of natural killer cells and other immune cells (30, 31). Due to its role in tumour-responsive innate immunity, it is a potential target in the development of anti-cancer therapies through modulation of NK cell-mediated cytotoxicity (32). Siglec-7 is a single-pass type 1 transmembrane protein known to bind α2,8-disialyl residues, found in the headgroups of multiple GSLs, including GD3, GT1b and GQ1b (33). Binding occurs in a solvent-exposed region of the distal N-terminus of the V-set Ig-like domain, which is the domain cloned and expressed here (34). Siglec-7 is also known to bind GD1a, through interaction with the α2,3-sialyl residue of GD1a via a secondary binding site in the same domain (33, 35). When initially incubated with giant liposomes containing 10% of each GSL in the presence of 50 mM NaCl (buffer 1), no binding could be measured for Siglec-7 to any GSL (Fig. 2a). The interaction between Siglec-7 and α2,8-disialyl residues relies on extensive hydrogen bonding (10 direct and 7 water-mediated) and one key salt bridge (34), so we trialled a salt-free hydration buffer (buffer 2) to minimise electrostatic screening. In this condition, siglec-7 binding was not only greatly enhanced but remained specific to the headgroups it is reported to bind (GD3, GT1b, GQ1b and GD1a) while no binding was detected for GM3 or GM1, supporting this is not a non-specific electrostatic binding effect (Fig. 2a). In addition, siglecs are known to be expressed on the cell surface in a multivalent manner and bind to clustered glycans on the target cell’s surface, thereby increasing the avidity (30). However here, the siglec-7 domain purified is monomeric, and the GSLs have apparent homogeneity (not clustered), which could explain the lack of measurable binding in the presence of salt. To oligomerise siglec-7, 50 µM NiCl_2_ was used to cluster the N-terminal his-tags (buffer 3), forming reversible high-order oligomers (36). The presence of NiCl_2_ rescues the binding of siglec-7 in buffer containing NaCl, supporting that protein oligomerisation is key to increasing the avidity of the interaction in the presence of increased ionic strength (Fig. 2b).

**Figure 2:**
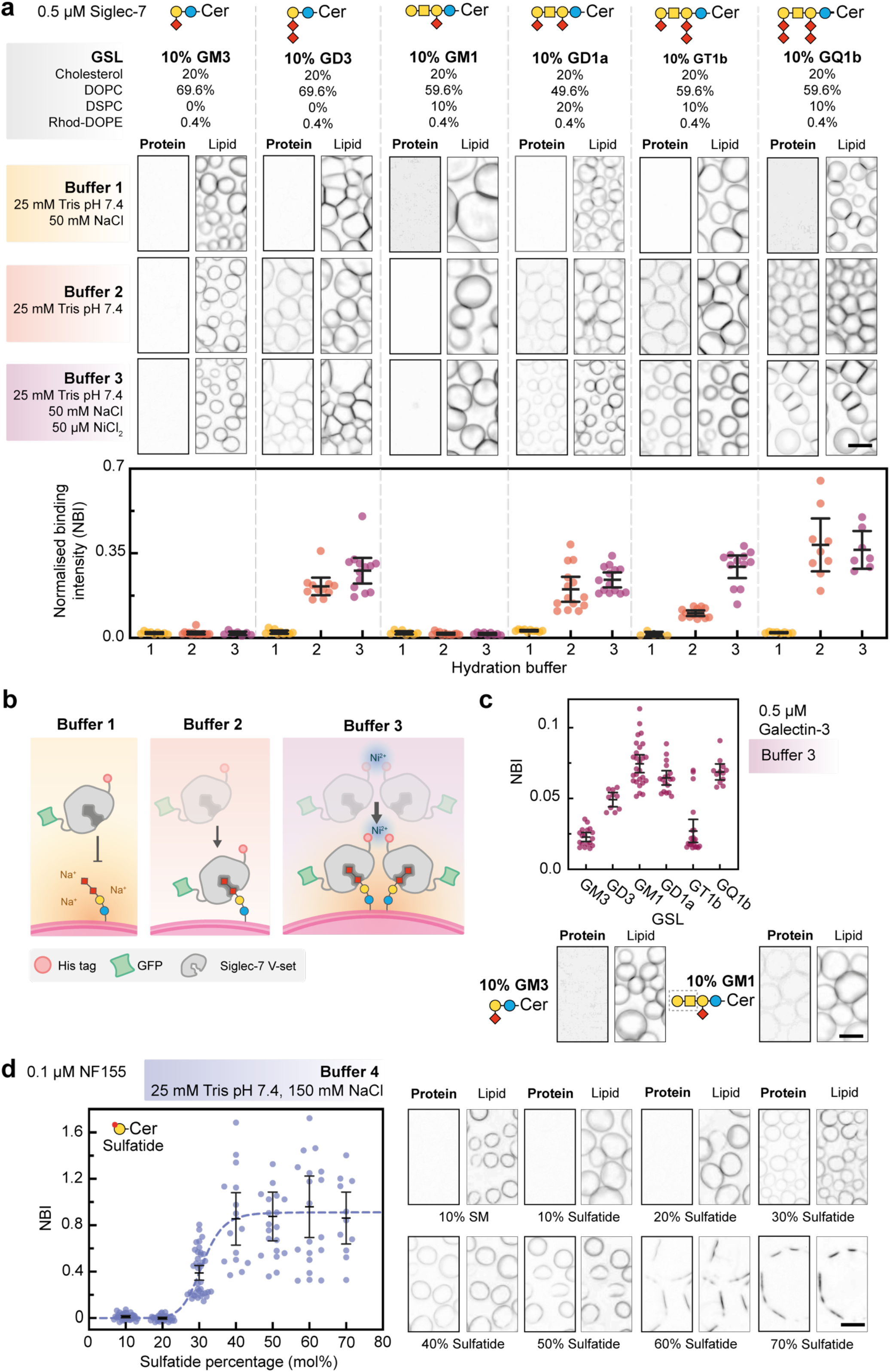
Using FLiPA to explore protein-GSL binding specificity and behaviour. **(a)** Siglec-7-GFP (0.5 µM) binding array to liposomes containing 10% GSLs, hydrated in either buffer 1 (25 mM Tris pH 7.4), buffer 2 (25 mM Tris pH 7.4, 50 mM NaCl) or buffer 3 (25 mM Tris pH 7.4, 50 mM NaCl, 50 µM NiCl_2_). Scale bar is 20 µm. *Below:* All individual image NBI values shown with mean ± 95% confidence interval. **(b)** Graphic representation of the influence of buffer composition and protein oligomerisation on siglec-7/GD3 binding. **(c)** Galectin-3-GFP (0.5 µM) binding array to liposomes containing 10% GSLs, hydrated in buffer 3 (25 mM Tris pH 7.4, 50 mM NaCl, 50 µM NiCl_2_). All individual image NBI values shown with mean ± 95% confidence interval. Scale bar is 20 µm. *Below:* Example images for GM3 and GM1 binding. Dashed box indicates Gal-3 binding ligand contained in GM1. **(d)** NF155-sulfatide binding varies with increasing sulfatide mol%. All individual image NBI values shown with mean ± 95% confidence interval and data fitted to the Hill equation. Example images for each sulfatide mol% are shown to the right. Scale bar is 20 µm.

The second bioreporter we utilised was galectin-3, a member of the galectin family of lectins (37). Galectin-3 is multi-functional and plays roles in a vast array of cellular processes including proliferation, differentiation and cell-cell interactions, has been found across a broad range of tissues such as immune, gastrointestinal and kidney cells, and has been shown to localise to the extracellular environment, the cytoplasm and the nucleus (38). It is formed of a carbohydrate recognition domain with a 113 amino acid-long N-terminal domain rich in proline and glycine residues. The N-terminal region lacks carbohydrate binding activity; however, it has been shown to participate with the CRD in glycan binding, is important for galectin-3 secretion, and has been demonstrated to be fundamental to galectin-3 oligomerisation through intra-and intermolecular hydrophobic interactions (39-42). For ease of purification however, only the CRD was expressed for this study. Implementing the buffers used for siglec-7, we found that galectin-3 also demonstrated no binding in the presence of 50 mM NaCl but could tolerate 50 mM NaCl in the presence of 50 µM NiCl_2_providing further evidence that the galectin-3 oligomerisation state is integral to biological function. The GSLs bound by galectin-3 were GM1 (most), GQ1b, GD1a, and GD3 (least), with no binding seen for GM3 or GT1b (Fig. 2c). This is different to the GSLs bound by siglec-7, demonstrating how FLiPA can be used to explore GSL-binding specificity. However, the magnitude of the NBI for galectin-3 is lower than siglec-7 indicating a weaker interaction. This is to be expected as galectin-3 specifically binds N-acetlylactosamine, which isn’t present in the GSLs tested here, and instead is found in the lacto-/neolacto-GSL series or on glycosylated proteins. Here, the GSLs tested contain the highly similar disaccharide of galactose linked to N-acetylgalactosamine, and while not an ideal substrate, galectin-3 has been reported to interact with GM1 and other GSLs to support clathrin-independent endocytosis, which our data support (43-45).

The third bioreporter we tested was neurofascin isoform 155 (NF155), a single-pass type I transmembrane protein found in the PM of myelin-producing cells (oligodendrocytes and Schwann cells) that possesses a large (115 kDa), highly glycosylated extracellular domain. NF155 is responsible for anchoring the myelin sheath to the axonal membrane at the paranode, which it does through interactions with the axonal PM proteins contactin-1 (CNTN1) and contactin associated protein (Caspr), and with GSLs, particularly sulfatide. The headgroup of sulfatide is a sulphated galactose, so compared with the other GSLs discussed thus far, it is relatively small, has a highly localised negative charge and can be incorporated into liposomes at much greater molar percentages (mol%). Previous work has demonstrated that the interaction between NF155 and sulfatide-containing liposomes relies on high local sulfatide concentration (45-55 mol%), with a sigmoidal binding mode, which is severely diminished once the protein is truncated (8). This suggests that NF155 interacts through multiple, cooperative sulfatide-binding sites and therefore relies on sulfatide clustering to increase avidity. When explored using FLiPA, sulfatide was readily incorporated into the giant liposomes over the range 10-50%. While some membrane was formed at 60-70%, these liposomes were less stable. Using FLiPA, the NBI was measured over a broad range of sulfatide concentrations (10-70 mol%) demonstrating that NF155 interacts with sulfatide-containing membranes in a cooperative manner and binds strongly even in the presence of high salt (buffer 4) and without the requirement for protein oligomerisation. This is consistent with NF155 containing multiple sulfatide-binding sites incorporating multivalency within the single, large extracellular region of this protein. The overall trend in these binding data were very similar to our previously described binding data using alternative approaches (8), but FLiPA was capable of detecting binding to sulfatide at a lower mol% (30%), suggesting that the sulfatide headgroups are more robustly displayed in the FLiPA set up.

By utilising three bioreporters with differential GSL-binding profiles and alternative avidity-enhancing molecular strategies, we have demonstrated how FLiPA can be used to determine specificity and explore lipid-concentration dependent effects, allowing details of membrane association behaviour to be revealed.

### FLiPA is a powerful platform for characterising lipid phase separation and protein-mediated GSL clustering

Within a bilayer, lipids seldom produce homogenous arrays, but instead laterally segregate into distinct phases. The dynamics of this segregation is mediated by both the biophysical properties of the lipids themselves and their complex interactions with associated proteins. Lipids undergo phase separation in response to different environmental cues, the primary being temperature (46). When the ambient temperature is above the lipid’s melting temperature (Tm), they have high lateral mobility within the membrane, and this phase is known as the liquid disordered phase (L_d_). For temperatures below their Tm, acyl-chain packing order increases and mobility decreases forming a solid (gel) phase (S_o_). If cholesterol is present, some lipids form an intermediate phase known as the liquid ordered phase (L_o_), which displays highly ordered acyl chains akin to the S_o_ phase but with the rapid mobility and rotational freedom of the L_d_ phase (47). There is growing support that in cellular membranes, nanometre-sized platforms of ordered lipids, with biophysical properties like the L_o_ phase, can form in conjunction with specific proteins to generate biologically active hubs involved in cell signalling, cell adhesion, protein trafficking and exo/endocytosis (48-50). These regions are commonly referred to as nanodomains, microdomains or lipid rafts and are enriched with cholesterol and sphingolipids, including the GSLs (23, 51, 52). Studying the existence of these domains in cells has been challenging and is complicated by the extreme protein and lipid complexity of cellular membranes. Simplified model systems represent a useful tool for exploring protein and lipid partitioning or clustering in dynamic, modifiable membrane environments. We therefore sort to explore how FLiPA can be used to visualise time-resolved dynamics of lipid phase separation and assess the contribution of specific proteins to this process.

To explore lipid phase separation using FLiPA, we began by using an established phase-separating lipid mixture containing equal amounts of cholesterol (33.2%), DOPC (33.2%) and DSPC (33.2%), with two lipid dyes, 0.2% Rhod-DPPE (1,2-dipalmitoyl-sn-glycero-3-phosphoethanolamine-N-(lissamine rhodamine B sulfonyl) and 0.2% NBD-DOPE (1,2-dioleoyl-sn-glycero-3-phosphoethanolamine-N-(7-nitro-2-1,3-benzoxadiazol-4-yl) (53). The Rhod-DPPE laterally segregates with the DSPC while the NBD-DOPE colocalises with DOPC upon phase separation. Using FLiPA, phase separation is seen on a large scale and domain coalescence can be tracked over time with ease due to the fixed nature of the giant liposomes (Fig. 3a/b, Supplementary Fig. 2a). In Fig. 3a, ordered domains form and coalesce over the space of 5 minutes which is clearly seen with a widefield microscope focused on the top plane of the liposome. This focus compromises edge resolution but as the liposomes are giant with minimal curvature, a large surface can be visualised. Over time the liposomes will fully separate into single L_o_ and L_d_ domains (Fig. 3b).

**Figure 3:**
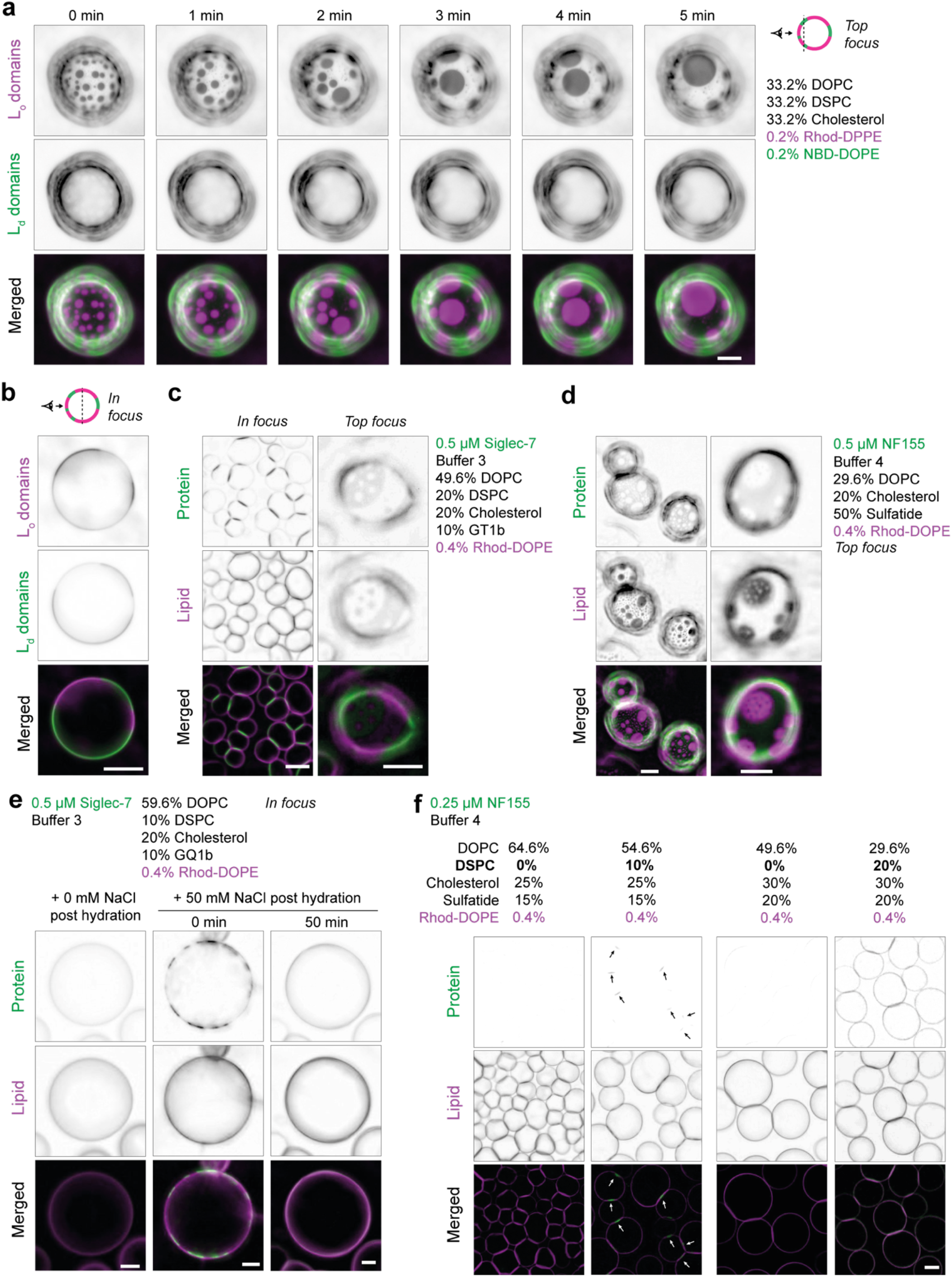
Using FLiPA to explore lipid phase separation and the effect of protein interactions. **(a)** Time resolved lipid-phase separation of a liposome containing 1:1:1 DOPC:DSPC:Cholesterol, with 2 distinct dyes that also phase separate, rhod-DPPE (ordered domains, magenta) and NBD-DOPE (disordered domains, green). Scale bar is 20 µm. **(b)** Phospholipid liposomes fully phase separate into 2 distinct ordered and disordered domains when allowed to equilibrate over time (2 hours). Scale bar is 20 µm. **(c)** Examples of lipid phase separation seen using 10% GT1b and 0.5 µM siglec-7-GFP. Scale bar is 20 µm. **(d)** Examples of lipid phase separation seen using 50% sulfatide and 0.5 µM NF155-GFP. Scale bar is 20 µm for the first example and 10 µm for the second example. **(e)** Osmotic pressure can be used to trigger phase separation of liposomes containing 10% GQ1b in the presence of 0.5 µM siglec-7-GFP by modulation of membrane tension, which dissipates as the aqueous environment across the membrane re-equilibrates. Scale bar is 10 µm. **(f)** Binding of 0.25 µM NF155-GFP can be enhanced by local enrichment of sulfatide using increased mol% of DSPC. Arrows highlight NF155/sulfatide microdomains. Scale bar is 20 µm. All images shown in this figure were taken at 20x magnification.

We next explored lipid phase separation in the presence of protein. As GSLs laterally segregate with ordered lipid they are found predominantly in the L_o_ phase, which is thought to underpin much of their biological function as GSL-rich nanodomains. Using Rhod-DOPE, the L_d_ domains can be visualised, and the addition of a GSL-binding GFP-fused protein will allow indirect localisation of the GSL-containing L_o_ domains. As mentioned previously, when 10% GT1b is added to lipid mix 4 (20% DSPC, 20% cholesterol), the membrane phase separates which can be visualised using Siglec-7-GFP (Fig. 3c). Interestingly, GSL and protein-dependent phase separation has distinct characteristics to phospholipids alone, for instance the GSLs are more commonly found at liposome-liposome contact sites and full phase separation is uncommon, with smaller L_o_ or L_d_ domains being sustained over time. This latter observation is pertinent in the context of the dynamics of lipid nanodomains *in vivo*, as proteins are thought to play an integral role in establishing and maintaining this compartmentalisation at physiological temperature (50). A similar behaviour is also seen in lipid mixtures that contain 50% sulfatide, incubated in the presence of NF155-GFP (Fig. 1d). In this case microdomains are seen within microdomains and some remain extremely small (although still micron-sized) over extended periods of time (24 hours). An important caveat with FLiPA is that it has been optimised to be used at 20x magnification, therefore smaller domains at the nanoscale could be forming but not resolved. Higher magnification analysis of sub-micron domains could be studied within the 96-well plate, or liposomes can be transferred to a glass slide for oil-immersion microscopy.

Another observation seen with GSL/siglec-7-GFP interactions is that in buffer 2 (0 mM NaCl) 0.5 µM siglec-7-GFP initially binds the membrane with apparent homogeneity but upon the addition of 50 mM NaCl, GFP signal localises to small, mobile puncta on the membrane surface which then dissipated back to apparent homogeneity over time (Fig. 3e). As mentioned, TAL-embedded liposomes are sensitive to pipetting so great care must be taken when adding NaCl to the buffer. This NaCl-dependent membrane reorganisation is likely caused by the application of external osmotic pressure, reducing the internal aqueous volume which in turn decreased membrane tension and facilitated phase transition. Over time membrane tension is restored as NaCl concentration across the membrane equilibrates, resulting in reestablishment of the original phase. We also observed this with NF155-GFP binding to liposomes containing 33.2% sulfatide, however the phase transition was sustained over time, indicating that some protein-GSL interactions stabilise ordered membrane phases (Supplementary Fig. 2b). The impact of osmotic pressure on lateral lipid arrangement has been described in detail for model membranes, particularly those containing phospholipids, but not for GSL-containing membranes. Our observation that GSL/protein phase separation occurs in response to change in osmotic pressure may have physiological relevance to cell function (54, 55). In support of this hypothesis, it was shown in yeast that increasing external salt concentration triggered spontaneous relocalisation of PtdIns(4,5)P_2_ into discrete regions within the PM, which inactivated the multiprotein signalling complex TORC2 (56).

As demonstrated here, many proteins that interact with GSL-rich nanodomains exploit different biological mechanisms to increase avidity. For NF155 this relies on local sulfatide concentration exceeding 30%, likely due to multiple low-affinity binding sites across the protein surface functioning together to promote binding (8). We hypothesised that we could explore NF155 binding at lower total sulfatide mol% (20% and below), which does not permit binding when the carrier lipid is DOPC alone (Fig. 2d), if we added DSPC to encourage lipid phase separation inducing local enrichment of sulfatide within L_o_ phases. Adding 10% DSPC and 25% cholesterol to liposomes containing 15% sulfatide, NF155 binding was seen, particularly at liposome contact sites (Fig. 3f, *left*). As many GSLs prevent formation of liposomes at total mol% above 10%, this demonstrates an alternative approach to increase the local GSL concentration by formation of L_o_ domains. This method was used previously to explore PtdIns(4,5)P_2_ -dependent protein crowding by locally increasing PtdIns(4,5)P_2_ concentration using DPPC (57). We also used a membrane composition containing 20% DSPC, 30% cholesterol and 20% sulfatide, and interestingly this increased NF155 binding in an apparently homogenous manner (Fig. 3f, *right*). This result suggests that nano-sized L_o_ domains may be forming even though they cannot be resolved as NF155 binding relies on clustering of the sulfatide headgroup.

Taken together, these observations demonstrate the considerable utility of FLiPA for exploring the dynamics of lipid phase separation and protein-dependent transition effects, offering a highly tuneable platform to further explore the roles of lateral lipid segregation in protein-GSL interactions.

## Discussion

Biological processes often require transient, spatially controlled recognition rather than maximally tight binding. Protein-GSL interactions are key examples of low-affinity/high-avidity interactions that provide physical and regulatory advantages over a single ultra-high-affinity interaction. Weak interactions allow reversible, dynamic remodelling of contacts but pose challenges for *in vitro* biophysical quantification, which FLiPA can overcome. Multivalency can be enhanced through clustering of GSLs in ordered membrane domains increasing the local concentration, oligomerisation of soluble proteins, clustering of membrane proteins and multiple GSL-binding sites contained within single proteins (Fig. 4). *In vitro* techniques need to reflect this complexity and often utilise model membranes, including micelles, liposomes and supported mono- and bilayers. It is widely accepted that membrane composition and properties influence protein-GSL interactions, with different model membranes yielding different affinities for the same protein-GSL interaction (58). As there are many variables that influence membrane formation, organisation and stability, FLiPA provides an optimisable toolkit for the robust characterisation of protein-GSL interactions, allowing comprehensive screening and side-by-side comparison of variables such as lipid composition and buffer components.

**Figure 4:**
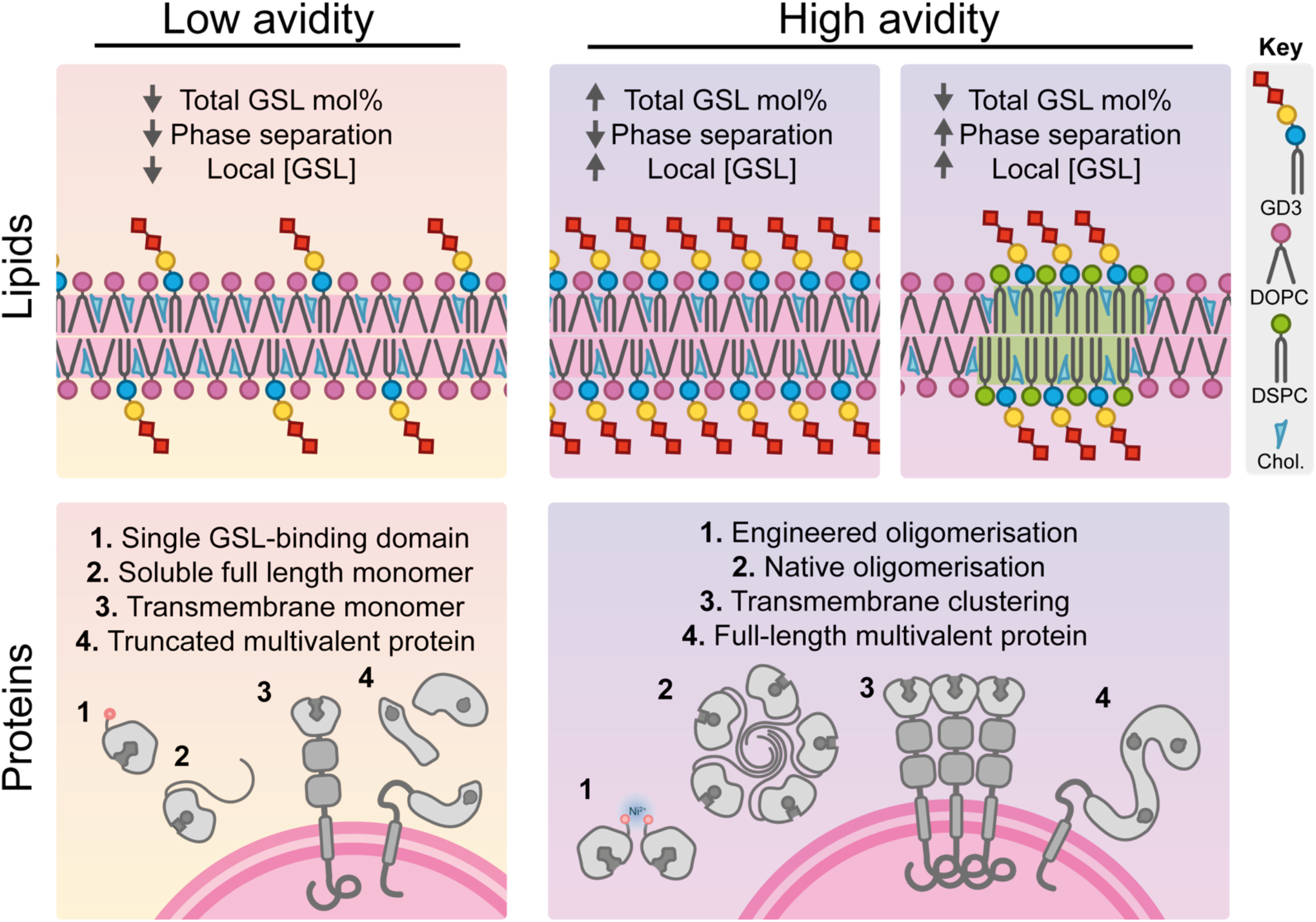
The strategies with which proteins and lipids increase avidity to enhance binding interactions and perform biological function.

We have demonstrated clearly that the ratio of saturated and unsaturated carrier lipids (DSPC:DOPC) governs the stable incorporation of GSLs in the bilayer, preventing aggregation and ensuring carbohydrate headgroups are optimally arrayed. The larger the GSL headgroup, the higher the mol% of DSPC was required to form giant liposomes. The importance of this observation is supported by previous work demonstrating that DSPC/GM1 liposomes bind the most strongly to the cholera toxin B-subunit, rather than DOPC/GM1 and DPPC/GM1, potentially reflecting better GSL incorporation and headgroup presentation when GM1 is incorporated into a more ordered membrane (59). This highlights another way in which FLiPA could be utilised to explore how membrane fluidity affects protein binding as varying the DSPC:DOPC ratio is straightforward.

Optimising GSL incorporation across a panel of 6 different GSLs facilitated the clear dissection of binding specificities for 2 GFP-fused bioreporters, siglec-7 and galectin-3. We found that in the presence of NaCl, oligomerising these bioreporters was crucial for detecting the interaction in FLiPA. The addition of NiCl_2_ offers a rapid solution for inducing oligomerisation of His-tagged proteins without the need for additional multimerisation domains that may influence protein folding or lead to steric hinderance during GSL-binding. Although siglec-7 and galectin-3 were also able to bind specific GSLs in the absence of NaCl, the oligomerisation of these proteins enhanced the affinity such that the electrostatic screening caused by the presence of NaCl could be overcome. This approach enables the use of physiologically relevant buffer systems for weak interactions and better reflects the biological behaviour of full length siglec-7 and galectin-3, which form transmembrane clusters and soluble oligomers, respectively (30, 39). Siglec-7 demonstrated binding to GSLs that contained α2,8-disialyl residues including GD3, GT1b and GQ1b as well as to the α2,3-sialyl residue of GD1a, all of which are supported using alternative methods (33, 35). Galectin-3 displayed weaker yet differential binding to the GSL panel, interacting with GM1, GD1a and GQ1b. The weak binding in comparison to siglec-7 is in line with what we know about galectin-3’s preferred binding substrate, N-acetlylactosamine, which is not present in the GSLs used here. However, previous work suggested it can bind GM1, which we corroborate, but also extend this to the more sialyated GD1a and GQ1b (43-45). Utilising a third bioreporter, the full-length extracellular domain of NF155, which is known to bind the GSL sulfatide, we were also able to demonstrate that FLiPA can be used to reveal further information on the way in which a protein interacts with the membrane. The sigmoidal binding response seen for NF155 as the mol% of sulfatide is increased supports the model that NF155 interacts with sulfatide via multiple low affinity sites to produce a high avidity interaction. NF155 does not require oligomerisation for binding and interacts strongly with the membrane in 150 mM NaCl. The use of these 3 diverse bioreporters demonstrates how powerful FLiPA is for determining the molecular details of GSL-protein interactions, allowing clear exploration of headgroup specificity, buffer sensitivity and membrane order. It also enables screening to identify new GSL-binding proteins and facilitates testing and enhancement of specific GSL binders that could be developed as new molecular probes in cells.

Despite ongoing debate regarding the lipid raft model, it is widely accepted that interactions between sphingolipids, cholesterol and proteins promote the formation of membrane domains in the outer leaflet of the PM that contribute to selective transport and signalling processes (51). Membrane compartmentalisation is thought to arise from a complex interplay between the biophysical properties of the lipids themselves and the cooperative effects of the proteins with which they interact. However, despite substantial research in this area, how these interactions support cellular function remain difficult to probe experimentally. FLiPA offers an additional tool to explore the properties of membrane phase separation, particularly in the context of GSL-containing membranes and how this is influenced by specific GSL-binding proteins. FLiPA benefits from excellent visual and temporal resolution of bilayer phase separation and exquisite control over membrane composition, protein oligomerisation and buffer constituents. We have identified specific lipid mixtures that support effective GSL presentation and demonstrated that FLiPA can reveal the dynamics of lipid phase separation. We also provide examples of how proteins (siglec-7 and NF155) can influence these dynamics, yielding small domains that do not coalesce over time. In addition, we have provided strategies for increasing the local GSL concentration through the formation of microdomains to enhance interactions with GSL-binding proteins that rely on avidity-enhancing mechanisms.

Further adding to the versatility of FLiPA, we have shown that membrane tension can be manipulated through producing osmotic pressure and that this leads to protein/GSL phase separation. This effect is well documented for mixtures of phospholipids, but characterisation is lacking for the GSLs (54). Cells have mechanisms to sense osmotic pressure through the change in membrane tension and respond accordingly to prevent cell lysis or shrinkage (54). Along with demonstrating that GSL phase separation occurs in response to osmotic pressure, we also revealed that the dynamics of this effect vary depending on the bioreporter/GSL used (Fig. 3e, Supplementary Fig. 2b), suggesting the interplay between specific proteins and lipids modulates the properties of the microdomains formed. This observation confirms that GSL phase separation occurs spontaneously after the application of osmotic pressure and this may influence protein activity, forming a mechanistic basis for sensing and responding to changes in membrane tension (55).

In summary, we have demonstrated that FLiPA is a highly versatile and informative *in vitro* platform for in-depth molecular characterisation of protein-lipid interactions and their impact on membrane behaviour. We have specifically designed this to be accessible for any biochemistry laboratory enabling broader exploration of these complex interactions. Our focus on the GSLs demonstrates how this approach can be fine-tuned to reveal mechanistic insight into the behaviour of these complex lipids and the proteins with which they interact, representing a significant advancement for *in vitro* methodologies in this field.

## Methods

### Reagents

Lipids used were purchased from Avanti with full details found in supplement. All lipids were stored at -20°C in glass vials under nitrogen gas and stable for 2-3 months. Coding sequences for GSL binding bioreporters were ordered from Thermofisher GeneArt, with NF155 and Siglec-7 codon optimised for human expression and Galectin-3 optimised for *Escherichia coli* expression (Supplement Table 1).

### Recombinant HEK293F protein expression

NF155 full length extracellular domain and Siglec-7 Ig-like V-type domain were cloned into the pHLSec plasmid with and N-terminal His_10_ tag and a C-terminal superfolder Green Fluorescent Protein (sfGFP) tag and secretion signal. The plasmids were transfected into 30 mL of Expi HEK293F cells (Expi293F™, A14527), cultured in Expi293™ Expression Medium (A1435101) to a density of 1.5 million cell/mL. Per 30 mL culture, 30 µg of plasmid DNA was mixed with 60 µg of PEI MAX® transfection reagent in 3 mL of Opti-MEM™ (31985062) and incubated for 20 min at room temperature before drop-wise addition to the Expi HEK293F culture. Cells were then incubated at 37°C with shaking for 4-5 days to allow for protein expression.

### Recombinant *E. coli* protein expression

Galectin-3 was cloned into the pOPTH plasmid with an N-terminal His_6_ tag and a C-terminal enhanced GFP (eGFP) tag. The plasmid was transformed into BL21(DE3) Competent E. coli (C2527), and the cells were cultured in 2YT media at 37°C until the OD_600_ reached 0.6. To induce protein expression, 0.5 mM IPTG was added, and the culture was incubated over night at 21°C. Cells were pelleted by centrifugation at 4000 x g using the Beckman Coulter JLA-8.1000 Fixed-Angle Rotor for 20 minutes at 4°C and resuspended in 30 mL 1xPBS before centrifugation again using the same conditions. The supernatant was removed and the pellet was resuspended in lysis buffer (20 mM Tris HCL pH 7.4, 500 mM NaCl, 0,5 mM TCEP, 20 mM imidazole and EDTA free protease inhibitors) before cell disruption under pressure. The lysed cells were centrifuged at 40, 000 x g in the Beckman Coulter JA-25.50 fixed-angle rotor for 30 minutes at 4°C and the supernatant was collected.

### Protein purification

For NF155-GFP and Siglec-7-GFP, proteins were harvested from 30 mL of HEK293-Expi cells by centrifugation at 100g for 5 minutes to pellet and remove the cells, followed by centrifugation of the resulting clarified media at 4000g to remove remaining debris. The fully clarified media was then sterile filtered through 0.2 micron Stericup filter units. The clarified media was incubated with NiNTA agarose beads (Qiagen, 30210) for 1 hour on a rolling bed at room temperature. The beads were transferred to a Bio-Rad Econo-Column^®^ (7374156) and separated from the media before being washed in wash buffer containing 25 mM Tris HCL pH 7.4, 150 mM NaCl and 10 mM imidazole. The bound protein was then eluted using elution buffer containing 25 mM Tris HCL pH 7.4, 150 mM NaCl and 300 mM imidazole. The eluted protein was concentrated to 1-2 mg/mL using Merck Amicon™ Ultra-15 Centrifugal Filter Units with a 30 kDa cutoff (UFC903024), dialysed overnight into storage buffer (25 mM Tris HCl, pH 7.4, 150 mM NaCl) and snap frozen in liquid nitrogen before long-term storage at -70°C.

For Galectin-3, the supernatant from the lysed BL21(DE3) *E. coli* was loaded onto a His-Trap™ FF 5 mL column (Cytiva, 17-5255-01) using the ÄKTA pure™ and washed using wash buffer (20 mM Tris HCl, pH 7.4, 500 mM NaCl, 20 mM imidazole) before elution in in elution buffer (20 mM Tris HCl, pH 7.4, 500 mM NaCl, 300 mM imidazole). The eluted fractions were pooled and further purified using the Superdex ™16/600 s75 gel filtration column, equilibrated in 50 mM Tris HCl, pH 7.4, 150 mM NaCl and 0.5 mM TCEP. The fractions containing purified protein were pooled, concentrated to 2.3 mg/mL using Merck Amicon™ Ultra-15 Centrifugal Filter Units with a 10 kDa cutoff (UFC801024) and snap frozen in liquid nitrogen before long-term storage at -70°C.

### FLiPA procedure

A 1% agarose solution was prepared using ultra-low gelling temperature Agarose (type IX) (Sigma, A2576-5G) in filtered water. Once fully melted, 50 µL of the 1% agarose mixture was pipetted into 12 wells of an organic-solvent resistant, glass-coated 96-well plate (Thermo Fisher, 60180-P334). Once the bottom of the 12 wells were fully coated and air bubbles removed using a needle, the 50 uL was fully aspirated from each, leaving behind a residual layer of agarose. This was repeated until the whole plate was coated. The ultra-thin agarose layer was dried fully either by incubation at room temperature overnight or placing on a 50°C hot plate for 2-3 hours. For long term storage, the wells were taped to prevent dust and moisture entering the wells and stored in a sealed desiccated bag and kept at room temperature.

Lipid mixtures were prepared in chloroform at 3 mM total lipid and then 3 µL was pipetted carefully onto the agarose layer in the well of the 96-well plate. The solvent was left to fully evaporate for at least 5-10 mins on a 50°C hotplate and the plate was used immediately. The giant liposomes were hydrated using a buffer containing 25 mM Tris-HCl, pH 7.4 and 0.1-0.5 µM GFP-tagged protein. Depending on the experiment, 50-150 mM NaCl and/or 50 µM NiCl_2_ was also present. Liposomes and GFP-tagged protein were incubated for 1 hour (unless otherwise stated) at room temperature before being imaged.

### Image acquisition and image processing

Images were taken using the EVOS M5000 imaging system fitted with a 3.2 MP monochrome CMOS camera (2,048 x 1,536 pixels) with 3.45 μm pixel resolution, and 2 long working distance objectives (10x and 20x). The GFP-tagged protein and Rhodamine-DOPE labelled membrane were imaged using the GFP (482/524 nm) and RFP (542/593 nm) light cubes, respectively. Light intensity and exposure length settings were fixed when images were compared within/between experiments for consistent quantification.

Image processing was performed using two open-source image analysis programs, CellProfiler (https://cellprofiler.org/) and FIJI (https://imagej.net/). For NBI measurement, we created a CellProfiler pipeline, emulating the data analysis performed by Saliba et al (27) (Supplementary Fig. 1a). We were unable to use the published pipeline as a number of modules were no longer supported by CellProfiler. For FLiPA, analysis began with converting the coloured images to grayscale and then filtering out background using the EnhanceOrSupressFeatures module. The image attained of the Rhod-PE-labelled membranes was used to identify regions of protein-lipid interaction, so a threshold filter was applied. Over-exposed pixels were identified in both the membrane and protein images and subtracted from the membrane threshold image. The resulting mask was applied to both the grayscale lipid and protein images after background subtraction and mean intensity was calculated for both and used to calculate NBI:

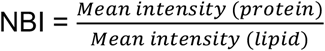

For visualisation of the images, FIJI was used to first convert images to 16-bit, followed by background subtraction. For separate channels, images were inverted and contrast enhanced. For merged images, after background subtraction a magenta LUT (look up table) was applied to the membrane image and a green LUT was applied to the protein image before the channels are merged and contrast was enhanced.

For measurement of the liposome diameter, FIJI was used to first convert the membrane images to 8-bit followed by background subtraction. A gaussian blur (sigma = 1) was used and then a threshold was applied using the Otsu method. The image was converted to a mask, the rings were closed and the holes filled. Joined liposomes were separated using the watershed function in FIJI and particles were analysed for area as long as they met size (50-infinity pixels) and circularity (0.75 and above) criteria. The area was then converted to equivalent circular diameter (ECD) (Supplementary Fig. 1b):

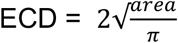

## Acknowledgments

We would like to thank Stephen Graham for gifting the his-tagged GFP, and Emma Weir and Henry Barrow for helpful discussions regarding microscopy image analysis. For the purpose of open access, the authors applied a Creative Commons Attribution (CC BY) license to any author accepted manuscript version arising from this submission.

## Funding

This work was supported by a Wellcome Trust Early Career Award (226903/Z/23/Z) to S.J.M, a Wellcome Trust Senior Research Fellowship (219447/Z/19/Z) to J.E.D., and E.B. is funded by the University of Cambridge School of Clinical Medicine Doctoral Training Programme in Medical Research.

## Contributions

S.J.M. contributed to research discussion, designed and optimised FLiPA, conducted the experiments and analysed the data. E.B. contributed to research discussion and design and performed cloning and protein purification of GSL bioreporters. J.E.D contributed to research discussion and design and provided technical expertise in GSL biology. S.J.M wrote the manuscript with support from both J.E.D and E.B.

## Notes

### Competing Interest Statement

The authors have declared no competing interest.

