## Supplemental Figures and Tables for "FLiPA: A versatile platform for quantitative analysis of protein-glycosphingolipid interactions"

This document includes:

Supplementary Figures 1-2
Supplementary Tables 1-2

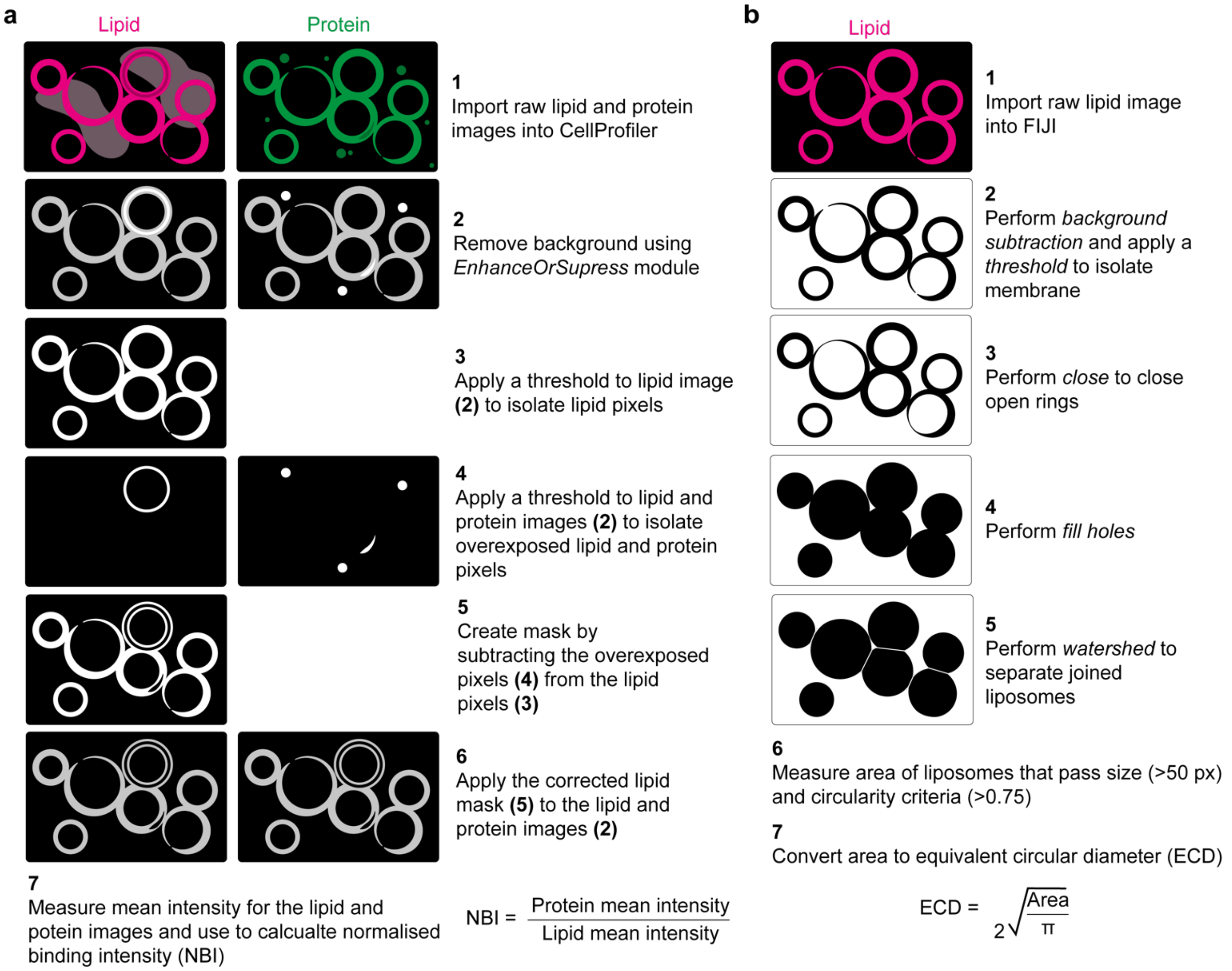


**Supplementary Figure 1:** Image processing procedures in FLiPA to determine liposome area, equivalent circular diameter and normalised binding intensity (NBI). **(a)** To measure liposome/GSL interactions, multiple images per condition are analysed using CellProfiler. Once images are converted to grayscale the background is removed using the EnhanceOrSupress module. A threshold is then applied to the lipid image to isolate pixels that represent membrane. A second threshold is applied to both the lipid and protein images to select overexposed pixels which are then subtracted from the membrane threshold to create a mask that is applied to the lipid and protein images and allows only the protein pixels that colocalise with membrane and are not overexposed to be measured. The mean intensity for the lipid and protein pixels is then calculated and used to calculate the NBI. **(b)** To measure liposome diameter, multiple images per condition are processed using FIJI. Once imported and converted to grayscale background subtraction is performed, and a threshold is applied to locate lipid membrane before converting image to a mask. Any incomplete liposome rings are closed, and the resulting holes are filed. The watershed function is used to separate any joined liposomes, and area is measured providing liposomes meet size (>50px) and circularity (0.75) criteria. Area is then converted to equivalent circular diameter.

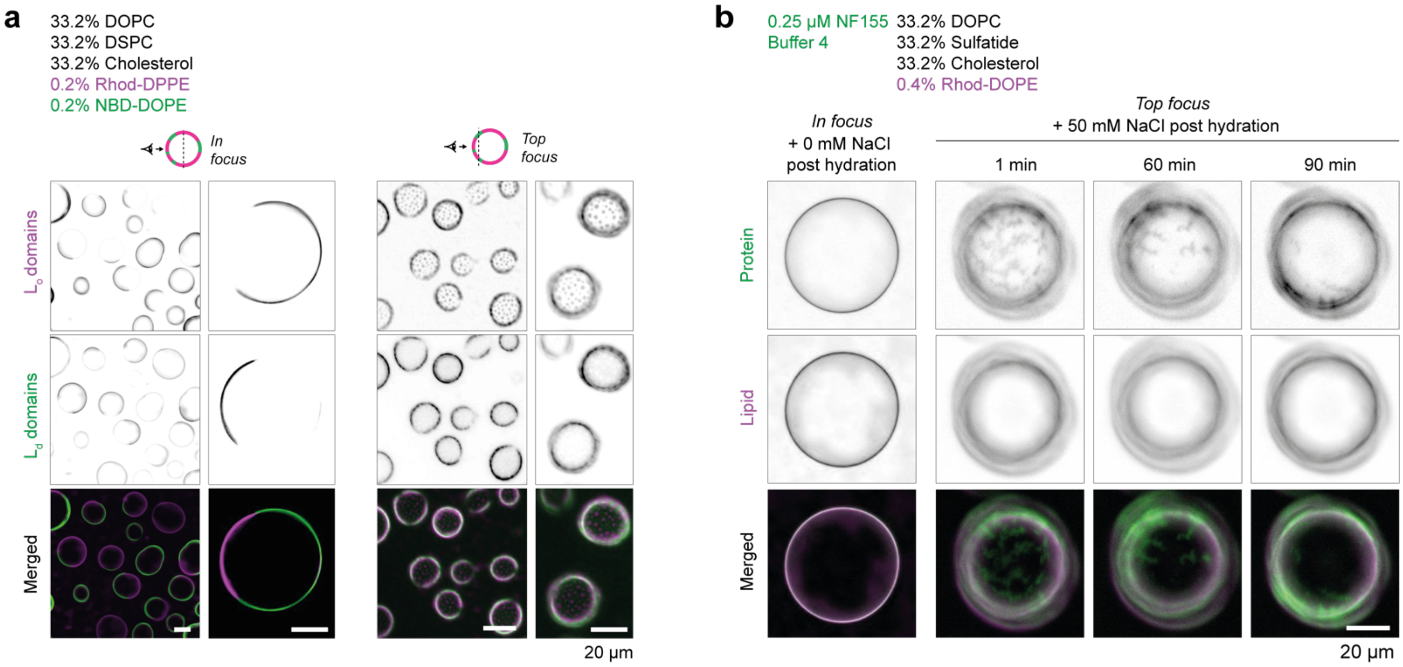


**Supplementary Figure 2:** Using FLiPA to explore the dynamics of lipid phase separation and the influence of GSL-binding proteins and osmotic pressure. **(a)** Further examples of lipid phase separation using FLiPA for a lipid mixture containing 33.2% DOPC, 33.2% DSPC, 33.2% cholesterol, 0.2% Rhod-DPPE (magenta, ordered domains) and 0.2% NBD-DOPE (green, disordered domains), viewed both in focus (later time point) and top focus (early time point). **(b)** Osmotic tension applied to sulfatide-containing liposomes labelled with NF155-GFP triggers phase separation which persists beyond 1 hour 30 minutes.

**Supplementary Table 1:** GSL binding bioreporters used to validate FLiPA.

| Protein name | Abbreviated name | Uniprot ID | Organism | GSL binding domain | GSL binding domain sequence | Fluorescent tag |
|---|---|---|---|---|---|---|
| Siglec-7 | Sig-7 | Q9Y286 | Human | Ig-likeV-type | QKSNRKDYSLTMQSSVTVQEGMCVHVRCSFSYPVDSQTDSDPVHGY<br>WFRAGNDISWKAPVATNNPAWAVQEETRDRFHLLGDPQTKNCTLSIR<br>DARMSDAGRYFFRMEKGNIKWNYKYDQLSVNVTA | sfGFP |
| Neurofascin isoform 155 | NF155 | O94856 | Human | NF155 extracellular domain (ECD) | IEIPMDLTQPPTITKQSAKDHIVDPRDNILIECEAKGNPAPSFHWTRNSR<br>FFNIAKDPRVSMRRRSGTLVIDFRSGGRPEEYEGEYQCFARNKFGTAL<br>SNRIRLQVSKSPLWPKENLDPVVVQEGAPLTLQCNPPPGLPSPVIFWM<br>SSSMEPITQDKRVSQGHNGDLYFSNVMLQDMQTDYSCNARFHFTHTI<br>QQKNPFTLKVLTNHPYNDSSLRNHPDMYSARGVAERTPSFMYPQGTA<br>SSQMVLRGMDLLLECIASGVPTPDIAWYKKGGDLPSDKAKFENFNKAL<br>RITNVSEEDSGEYFCLASNKMGSIRHTISVRVKAAPYWLDEPKNLILAP<br>GEDGRLVCRANGNPKPTVQWMVNGEPLQSAPPNPNREVAGDTIIFRD<br>TQISSRAVYQCNTSNEHGYLLANAFVSVLDVPPRMLSPRNQLIRVILYN<br>RTRLDCPFFGSPIPTLRWFKNGQGSNLDGGNYHVYENGSLEIKMIRKE<br>DQGIYTCVATNILGKAENQVRLEVKDPTRIYRMPEDQVARRGTTVQLE<br>CRVKHDPSLKLTVSWLKDDEPLYIGNRMKKEDDSLTIFGVAERDQGSY<br>TCVASTELDQDLAKAYLTVLGRPDRPRDLELTDLAERSVRLTWIPGDAN<br>NSPITDYVVQFEEDQFQPGVWHDHSKYPGSVNSAVLRLSPYVNYQFR<br>VIAINEVGSSHPSLPSERYRTSGAPPESNPGDVKGEGTRKNNMEITWT<br>PMNATSAFGPNLRYIVKWRRRETREAWNNVTVWGSRYVVGQTPVYV<br>PYEIRVQAENDFGKGPEPESVIGYSGEDYPRAAPTEVKVRVMNSTAISL<br>QWNRVYSDTVQGQLREYRAYYWRESSLLKNLWVSQKRQQASFPGDR<br>LRGVVSRLFPYSNYKLEMVVVNGRGDGPRSETKEFTTPEGVPSAPRR<br>FRVRQPNLETINLEWDHPEHPNGIMIGYTLKYVAFNGTKVGKQIVENFS<br>PNQTKFTVQRTDPVSRYRFTLSARTQVGSGEAVTEESPAPPNEATPTA<br>AYTNNQADIATQG | sfGFP |
| Galectin-3 | Gal-3 | P17931 | Human | Carbohydrate recognition domain (CRD) | LIVPYNLPLPGGVVPRMLITILGTVKPNANRIALDFQRGNDVAFHFNPRF<br>NENNRRVIVCNTKLDNNWGREERQSVFPFESGKPFKIQVLVEPDHFKV<br>AVNDAHLLQYNHRVKKLNEISKLGISGDIDLTSASYTMITGS | eGFP |

**Supplementary Table 2:** Lipids used to explore protein-GSL interactions and lateral membrane organisation in FLiPA.

| Abbreviated name | Systematic name | Lipid shorthand notation | Supplier | SKU | [stock] (mg/mL) | MW | [stock] mM | Solvent (ratio) |
|---|---|---|---|---|---|---|---|---|
| Liss Rhod DOPE | 1,2-dioleoyl-sn-glycero-3-phosphoethanolamine-N-(lissamine rhodamine B sulfonyl) | 18:1 | Avanti | A81150 | 1 | 1301.715 | 0.8 | chloroform |
| NBD DOPE | 1,2-dioleoyl-sn-glycero-3-phosphoethanolamine-N-(7-nitro-2-1,3-benzoxadiazol-4-yl) | 18:1 | Avanti | A81145 | 1 | 924.155 | 1.1 | chloroform |
| Liss Rhod DPPE | 1,2-dipalmitoyl-sn-glycero-3-phosphoethanolamine-N-(lissamine rhodamine B sulfonyl) | 16:0 | Avanti | A81158 | 1 | 1249.641 | 0.8 | chloroform |
| DOPC | 1,2-dioleoyl-sn-glycero-3-phosphocholine | 18:1 | Avanti | A80375 | 10 | 786.113 | 12.7 | chloroform |
| DSPC | 1,2-distearoyl-sn-glycero-3-phosphocholine | 18:0 | Avanti | A80365 | 10 | 790.145 | 12.7 | chloroform |
| DGS-NTA(Ni) | 1,2-dioleoyl-sn-glycero-3-[(N-(5-amino-1-carboxypentyl)iminodiacetic acid)succinyl] (nickel salt) | 18:1 | Avanti | A89404 | 5 | 1057.003 | 4.7 | chloroform |
| GM3 | Ganglioside GM3 | mixed | Avanti | A86058 | 5 | 1268.653 | 3.9 | (1:1) chloroform:MeOH |
| GD3 | Ganglioside GD3 | mixed | Avanti | A86060 | 5 | 1506.805 | 3.3 | (1:1) chloroform:MeOH |
| GM1 | Ganglioside GM1 | mixed | Avanti | A86065 | 5 | 1568.805 | 3.2 | (1:1) chloroform:MeOH |
| GD1a | Ganglioside GD1a | mixed | Avanti | A86055 | 5 | 1872.14 | 2.7 | (1:1) chloroform:MeOH |
| GT1b | Ganglioside GT1b | mixed | Avanti | A86059 | 5 | 2180.42 | 2.3 | (1:1) chloroform:MeOh |
| GQ1b | Ganglioside GQ1b | mixed | Avanti | A86086 | 2.5 | 2488.71 | 1.0 | (1:1) chloroform:MeOH |
| Sulfatide | 3-O-sulfogalactosylceramide | mixed | Avanti | A83305 | 5 | 897.857 | 5.6 | (2:1) chloroform:MeOH |
| Cholesterol | (3β)-cholest-5-en-3-ol | n/a | Merck | C8667 | 5 | 386.65 | 12.9 | Chloroform |
